# Mating promotes lactic-acid gut bacteria in a gift-giving insect

**DOI:** 10.1101/071001

**Authors:** Chad C. Smith, Robert B. Srygley, Emma I. Dietrich, Ulrich G. Mueller

## Abstract

Mating is a ubiquitous social interaction with the potential to influence the microbiome by facilitating transmission, modifying host physiology, and in species where males donate nuptial gifts to females, altering diet. We manipulated mating and nuptial gift consumption in two insects that differ in nuptial gift size, the Mormon cricket *Anabrus simplex* and the decorated cricket *Gryllodes sigillatus*, with the expectation that larger gifts are more likely to affect the gut microbiome. Surprisingly, mating, but not nuptial gift consumption, affected bacterial community structure, and only in Mormon crickets. The change in structure was due to a precipitous drop in the abundance of lactic-acid bacteria in unmated females, a taxon known for their beneficial effects on nutrition and immunity. Mating did not affect phenoloxidase or lysozyme-like antibacterial activity in either species, suggesting that any physiological response to mating on host-microbe interactions is decoupled from the systemic immunity. Protein supplementation also did not affect the gut microbiome in decorated crickets, suggesting that insensitivity of gut microbes to dietary protein could contribute to the lack of an effect of nuptial gift consumption. Our study provides experimental evidence that sexual interactions can affect the microbiome and suggests mating can promote beneficial gut bacteria.

---

Social interaction (Archie and Tung, 2015; Smith and Mueller, 2015) and diet (Ley *et al.*, 2008; Muegge *et al.*, 2011; Yatsunenko *et al.*, 2012; David *et al.*, 2014) are two key factors that influence the composition of the microbiome. Of the types of social interactions animals engage in, mating is both ubiquitous and among the most likely to influence host microbial communities due to its intimacy and profound effects on host physiology. Yet scant attention has been paid to the influence of mating on microbial symbiosis beyond the transmission of pathogenic infections (Lockhart *et al.*, 1996; Knell and Webberley, 2004) despite the fact that beneficial microbes can also be sexually transmitted during the mating process (Smith and Mueller, 2015). Mating also alters the expression of hundreds of genes involved in metabolism, reproduction, and immunity (McGraw *et al.*, 2008), which potentially could influence host-microbe interactions. The host immune system in particular plays a critical role in the regulation of the microbiome (Ryu *et al.*, 2010; Hooper *et al.*, 2012; Engel and Moran, 2013), which in turn influences host immune function (Hooper *et al.*, 2012; Engel and Moran, 2013; Levy *et al.*, 2015), nutrition (Turnbaugh *et al.*, 2006; Engel and Moran, 2013) and behavior (Archie and Theis, 2011; Forsythe and Kunze, 2013).

Sexual interactions can also influence diet, an important determinant of the constitution of the microbiome (Ley *et al.*, 2008; Muegge *et al.*, 2011; Yatsunenko *et al.*, 2012; David *et al.*, 2014). In many animals, males provide nuptial gifts that females ingest during courtship or copulation (Yosef and Pinshow, 1989; Vahed, 1998; Gomes and Boesch, 2009). Male crickets and katydids in particular are known for the production of a spermatophylax, a proteinaceous (Heller *et al.*, 1998), sperm-free mass that is eaten by females. Consumption of the spermatophylax has varying effects on female fitness, increasing survival and fecundity in some taxa (Gwynne, 1984a, 2008; Simmons, 1990) while producing no apparent benefit in other taxa (Will and Sakaluk, 1994; Vahed, 2007). This has led to extensive debate over spermatophylax evolution. Several lines of evidence suggest that the spermatophylax serves only as an ejaculate protection device to prevent the female from eating the sperm-laden ampulla (Vahed, 2007), which is transferred with the spermatophylax to females during copulation. These nuptial gifts are not necessarily expected to provide a nutritional benefit, only properties that distract the female long enough for sperm transfer to complete (Vahed, 2007). In contrast, the spermatophylax is expected to be nutritious when it serves as a form of paternal investment that increases the number or quality of offspring sired by the male (Gwynne, 2008). Which of these two explanations is correct is likely to have important implications for how nuptial gifts influence the microbiome, as protein intake can induce rapid changes in the gut microbial communities (Wu *et al.*, 2011; David *et al.*, 2014).

We manipulated nuptial feeding and mating to measure their effects on the gut microbiome of two insects that differ in the size of their gifts, the Mormon cricket, *Anabrus simplex* (Orthoptera: Tettiginiidae), and the decorated cricket, *Gryllodes sigillatus* (Orthoptera: Gryllidae). Mormon crickets produce a spermatophore six times larger than *G. sigillatus* (19% vs 3% of male body mass; Gwynne, 1984b; Sakaluk, 1985, (Fig. 1) and are a well-known example of nutrition-dependent sex-role reversal, with females competing for access to spermatophylax-producing males when food is scarce (Gwynne, 1984b, 1993). In contrast, the *G. sigillatus* spermatophylax is no larger than that required for sperm transfer (Sakaluk, 1984) and does not provide any detectable nutritional benefit to females (Will and Sakaluk, 1994; but see Ivy *et al.*, 1999). Given this evidence, we expect that spermatophylax consumption will exert larger effects on the gut microbiome of Mormon crickets than decorated crickets. Whether mating influences the microbiome depends on the potential for microbial transmission, as well as an effect of mating on the physiological state of females. We assessed these alternatives by screening male and female reproductive tissues for bacteria and measuring components of the immune system that are known to change in response to mating in insects.

**Figure 1.**
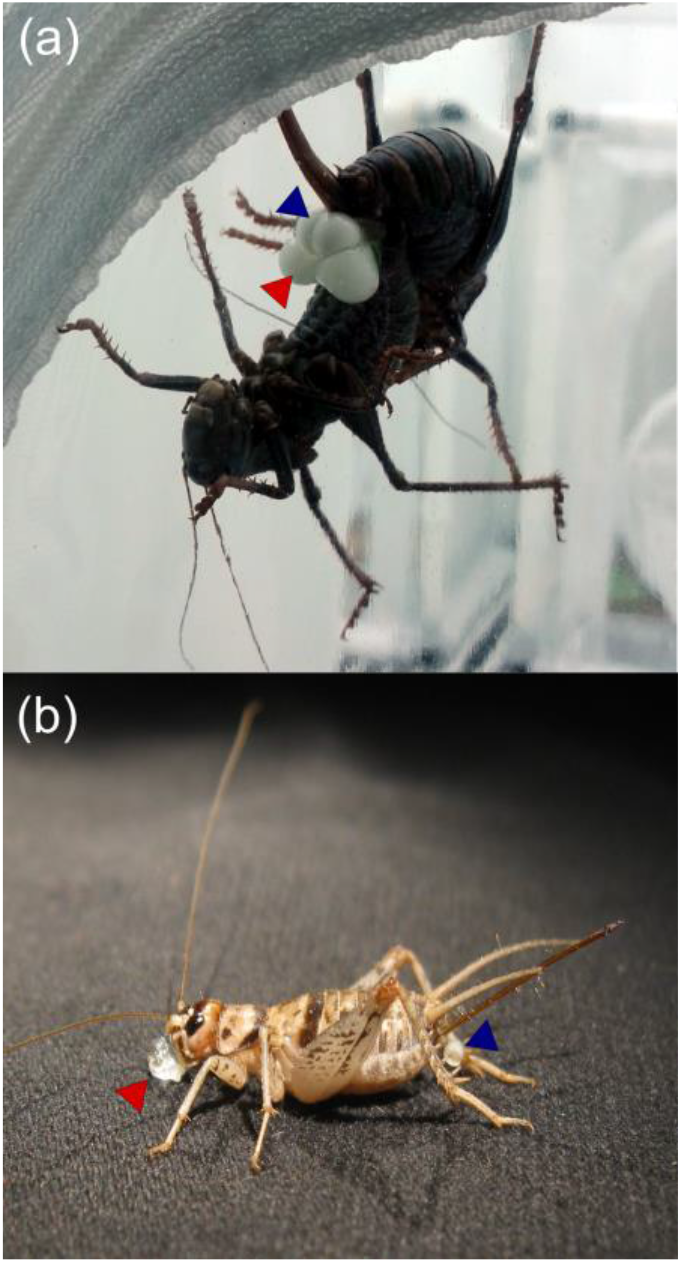
(a) Mormon cricket *Anabrus simplex* female (top) and male (bottom) in copula and (b) a decorated cricket *Gryllodes sigillatus* female after mating. Red arrow indicates the spermatophylax and blue arrow indicates the ampulla.

## RESULTS AND DISCUSSION

### Mating and the microbiome

We found that mating, but not spermatophylax consumption, influenced the structure of the gut microbiome of Mormon crickets (Figure 2, Table 1), while neither had an effect in decorated crickets (Table S1). Ordination of the Mormon cricket OTU scores suggested that five taxa changed in abundance in response to the mating treatment (Fig. 3), all lactic-acid bacteria (Family *Lactobacillaceae)*. Two of these were among the dominant members of the Mormon cricket microbiome (Figure S1, *Pediococcus acidilactici* 102222 and *Pediococcus sp*. 17309), while the other three occurred at a lower frequency *(Lactobacillus sp*. 288584, *Pediococcus sp*. 733251, and *Lactobacillus sp*. 1110317).

**Figure 2.**
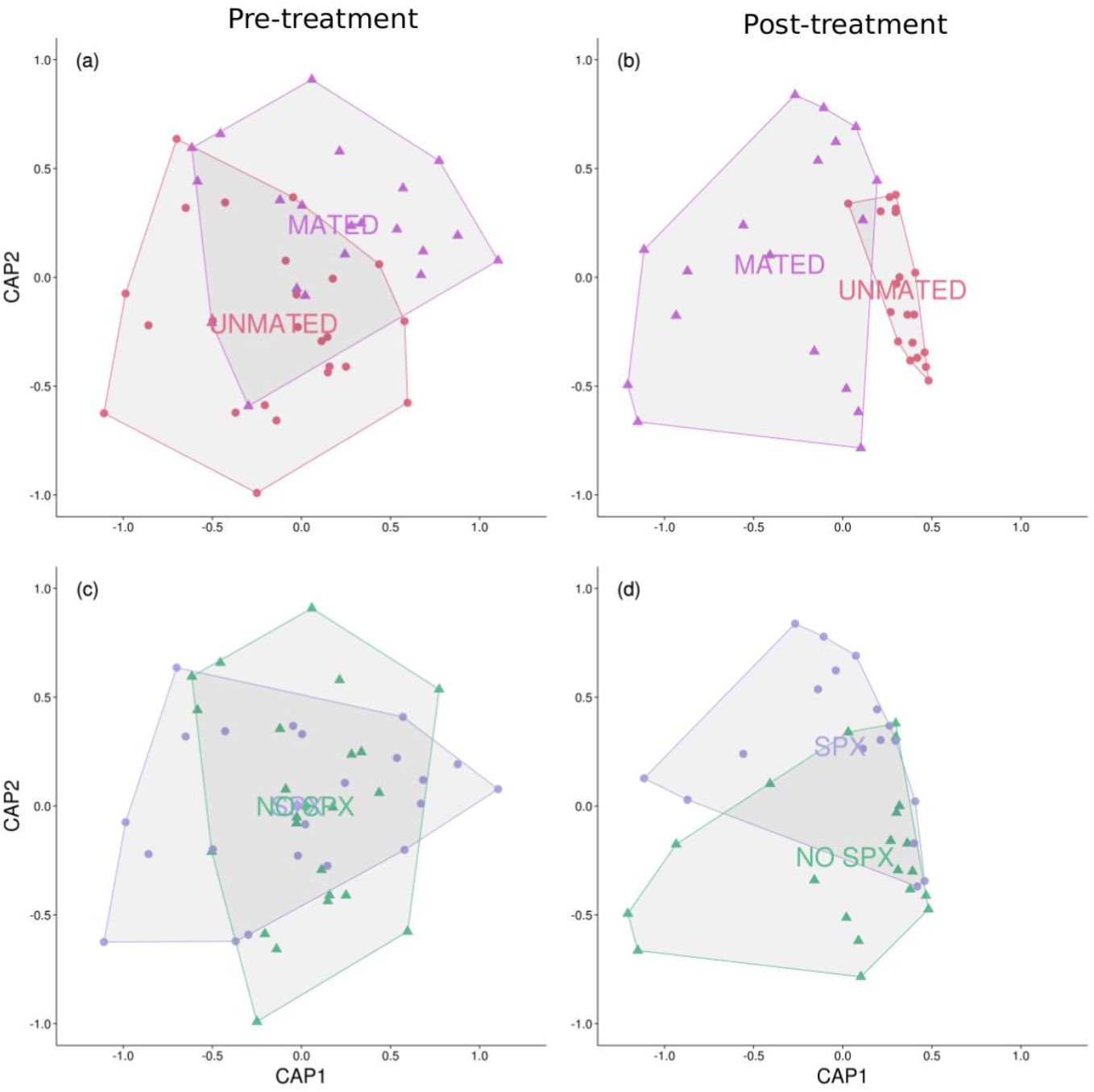
Ordination of Mormon cricket sample scores from a distance-based redundancy analysis. Points are colored to indicate whether a cricket was mated (triangles) or unmated (circles) (a,b) and whether they were allowed to consume the spermatophylax (circles) or not (triangles) (c,d). Text corresponds to the centroids for samples collected before (a,c) or after (b,d) the treatments were applied. Alpha diversity was not affected by mating or spermatophylax consumption (Table S2 and S3).

**Figure 3.**
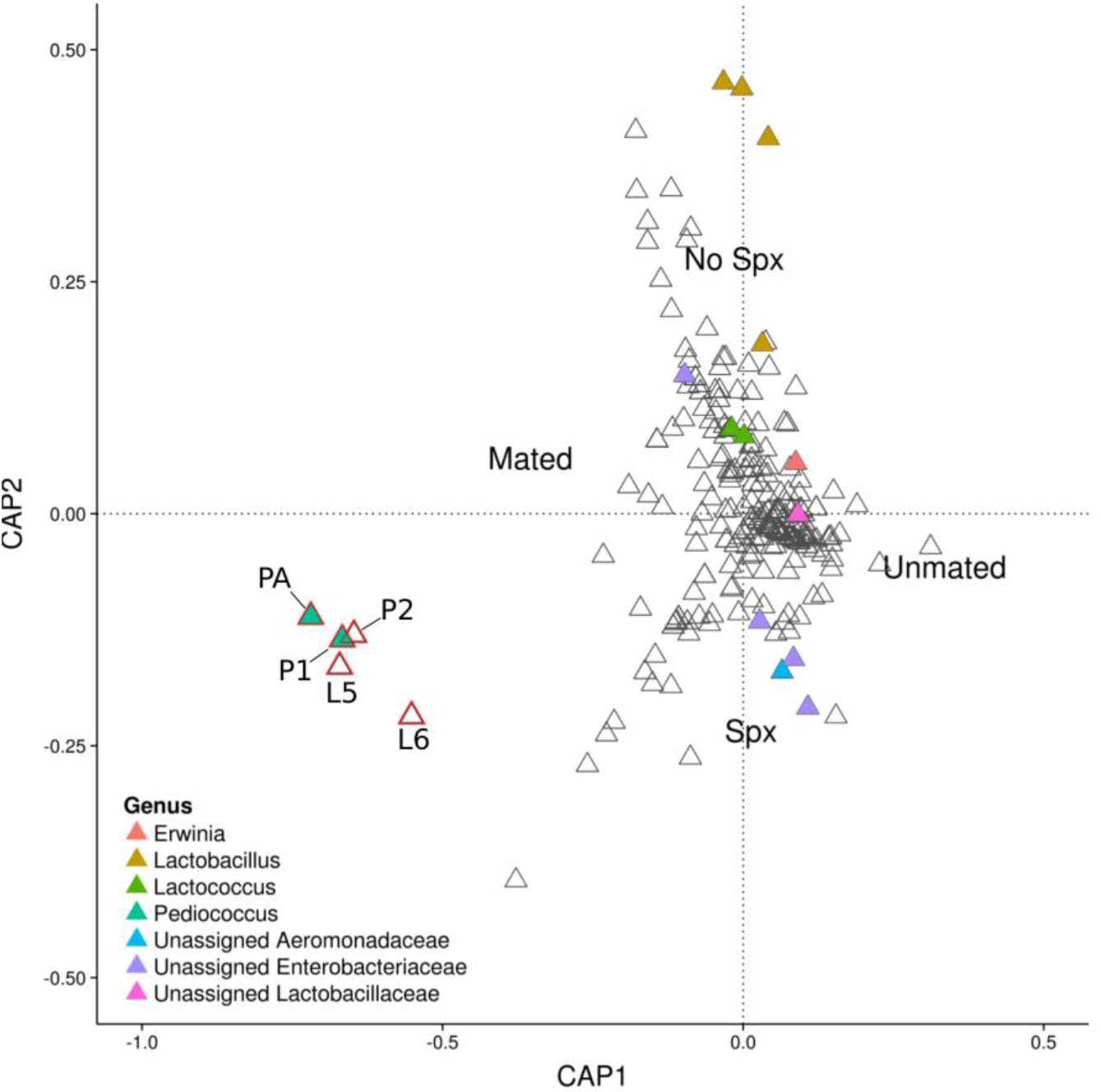
Ordination of Mormon cricket OTU scores from a distance-based redundancy analysis. Each triangle represents an OTU, with text indicating the centroid of the sample scores from each treatment. Filled triangles are the top 15 most abundant OTUs colored by genus. Labeled OTUs are those displaced along the axis associated with mating and individually analyzed for differences in abundance (see Figure 4, (Table 2). PA = *Pediococcus acidilactici* 102222, P1=*Pediococcus* 17309, P2=*Pediococcus* 773251, L5=*Lactobacillus* 288584, L6= *Lactobacillus* 1110317.

**Table 1.** Permutation tests from distance-based redundancy analysis of female Mormon cricket fecal samples. Time refers to whether a sample was collected before or after the treatments were applied.

|  |  | *F* | *P* |
| --- | --- | --- | --- |
| *Full model* | Mate | 0.67 | 0.66 |
|  | Spermatophylax | 0.19 | 0.99 |
|  | **Time** | **3.00** | **0.01** |
|  | Mate * Spermatophylax | 1.43 | 0.20 |
|  | Spermatophylax * Time | 0.61 | 0.71 |
|  | **Mate \* Time** | **2.39** | **0.04** |
| *Pre-experiment* | Mate | 0.76 | 0.74 |
|  | Spermatophylax | 0.38 | 0.99 |
|  | Mate * Spermatophylax | 0.89 | 0.55 |
| *Post-experiment* | **Mate** | **1.61** | **0.02** |
|  | Spermatophylax | 1.05 | 0.32 |
|  | Mate * Spermatophylax | 0.39 | 0.41 |

We compared the abundance of these five lactic-acid bacteria among treatments in univariate analyses and found that three differed depending upon whether females had mated or not, including *P. acidilactici* 102222 and *Pediococcus sp*. 17309 (Fig. 4, Table 2). Comparisons of fecal samples taken before and after the treatments indicated that all three lactic-acid bacteria experienced a precipitous decline in unmated females, but persisted in mated females, resulting in higher abundances in mated females at the end of the experiment (Fig. 4, Table 2).

**Figure 4.**
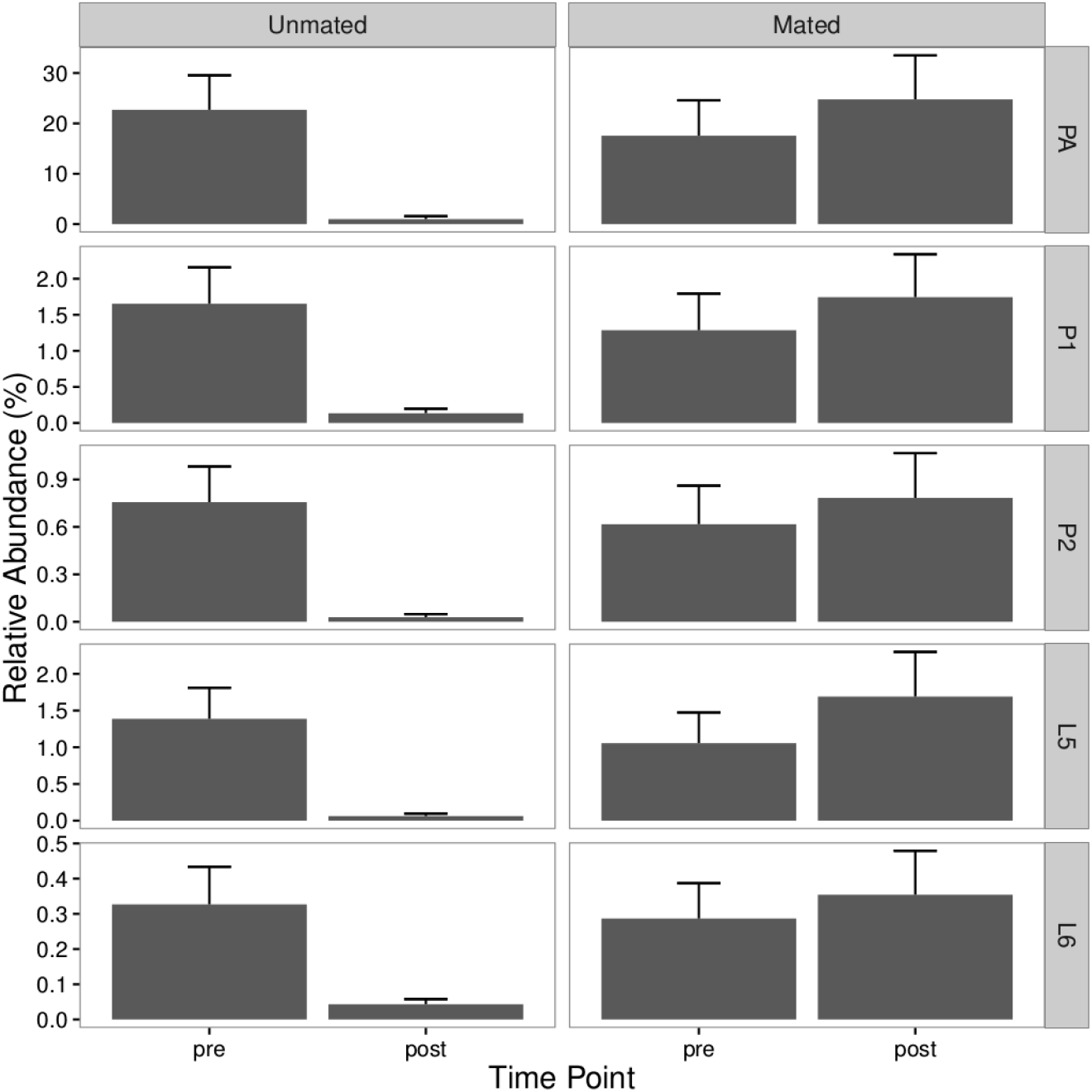
Relative abundance of five OTUs putatively associated with mating in Mormon crickets. Time point indicates whether samples were collected before or after the treatments were imposed. A significant interaction between mating and time point was detected for the top 3 panels (Table 2).

**Table 2.** OTU abundance of five taxa putatively associated with mating in Mormon crickets. Values represent the X^2^ (p-value) from an analysis of deviance, except for L6, which was analyzed with Wilcoxon signed-rank tests. Significant terms are in bold (p<0.05). PA = *Pediococcus acidilactici* 102222, P1=*Pediococcus* 17309, P2=*Pediococcus* 773251, L5=*Lactobacillus* 288584, L6=*Lactobacillus* 1110317.

|  |  | PA | P1 | P2 | L5 | L6 |
| --- | --- | --- | --- | --- | --- | --- |
| GLM | Mate | 3.26 (0.28) | 3.26 (0.28) | 3.01 (0.28) | 2.38 (0.28) |  |
|  | Spermatophylax | 0.24 (0.84) | 0.39 (0.82) | 0.25 (0.83) | 0.17 (0.85) |  |
|  | Time | **13.9 (0.006)** | 3.16 (0.28) | 3.14 (0.28) | 2.42 (0.28) |  |
|  | Mate * Spermatophylax | 0.01 (0.99) | 0.52 (0.77) | 0.02 (0.96) | 0.07 (0.92) |  |
|  | Spermatophylax*Time | 0.72 (0.68) | 2.42 (0.28) | 0.34 (0.83) | 0.05 (0.92) |  |
|  | Mate * Time | **12.1 (0.007)** | **10.7 (0.01)** | **8.30 (0.03)** | 5.20 (0.13)<sup>†</sup> |  |
| Wilcoxon | Pre-experiment samples: Mated vs. unmated females |  |  |  |  | 300 (0.85) |
|  | Post-experiment samples: Mated vs. unmated females |  |  |  |  | 100 (0.40) |
|  | Unmated females only: pre vs. post experiment |  |  |  |  | 200 (0.40) |
|  | Mated females only: pre vs. post experiment |  |  |  |  | 200 (0.99) |

<sup>†</sup> $P = 0.02$ before FDR correction for multiple tests.

Lactic-acid bacteria are known for their beneficial associations with the gastrointestinal tract of human and non-human animals (De Vos *et al.*, 2009; Walter *et al.*, 2011; Ma *et al.*, 2012), including insects (Forsgren *et al.*, 2010; Storelli *et al.*, 2011; Vásquez *et al.*, 2012; Erkosar *et al.*, 2015). *P. acidilactici*, for example, has been shown to enhance development and immune function (Neissi *et al.*, 2013), reduce susceptibility to infection (Castex *et al.*, 2009), and produce bacteriocins toxic to food-borne pathogens (Bhunia *et al.*, 1988). Our study thus shows that sexual interactions can influence the structure of the microbiome, and suggests that mating can promote the persistence of beneficial bacteria in the gut.

One way social behavior can alter the microbiome is by facilitating transmission of microbes between members of the group (Archie and Tung, 2015). Sexual transmission is unlikely to explain our results, however, because the male spermatophore and female spermatheca were negative in our 16s PCR screens for bacteria, perhaps because of antimicrobial activity in the reproductive tissues. Sexual transmission of both pathogenic and beneficial microbes, however, does occur in insects (Knell and Webberley, 2004; Smith and Mueller, 2015), and more studies are needed to evaluate their prevalence and effects on host fitness and reproductive behavior. Contact with male feces might also have provided a source of lactic acid bacteria to mated females if there are gender differences in the microbiome. While there is some evidence that gender influences the microbiome in other animals (Bolnick *et al.*, 2014; Ding and Schloss, 2014), this has yet to be evaluated in Mormon crickets.

Changes in host physiology in response to social interaction, or lack thereof, could also explain shifts in microbiome structure. Hormones that regulate appetite, energy expenditure, and metabolism are thought to affect the gut microbiome by altering (i) immune function, (ii) mucous production in the gut epithelia, and (iii) behavioral changes in food intake (Spor *et al.*, 2011). Similarly, the stress response (Jašarević *et al.*, 2015; Sandrini *et al.*, 2015) and fluctuations in reproductive hormones (Gajer *et al.*, 2012; Brotman *et al.*, 2014) are associated with changes in the composition of the microbiome. In *Drosophila*, mating influences the expression of >1700 genes involved in these physiological processes (McGraw *et al.*, 2008). Many of these genes are expressed in tissues outside of the female reproductive tract and are induced by the transfer of specific male seminal fluid proteins (McGraw *et al.*, 2008). Whether similar physiological responses to mating can be generalized to other insects, and whether these specific changes do influence host-microbe interactions, remains to be elucidated.

Mating in insects can result in the suppression of the immune system due to tradeoffs between survival and reproduction (Harshman and Zera, 2007), and the immune system is a key regulator of the microbiome (Ryu *et al.*, 2010; Hooper *et al.*, 2012). We measured three components of systemic immunity in both species and found that immunological activity was unaffected by mating, and was not associated with variation among crickets in microbiome structure (Table S4 and S5). This suggests that if the lactic-acid bacteria identified in our study are influenced by the immune system, it likely occurs locally within the gut rather than in response to systemic changes in immunity. This is consistent with experiments in *Drosophila*, where the immune response in gut epithelia is induced by oral introduction of bacteria but not after injection of the same bacteria into the hemocoel (Tzou *et al.*, 2000).

### Nuptial gift consumption and the microbiome

In contrast to our expectation that larger nuptial gifts should elicit a greater change in microbiome composition, spermatophylax consumption did not affect the gut bacterial communities in either species (Table 1, S1). At least three non-mutually exclusive possibilities could explain this result. First, it is possible that the spermatophylax is not a highly nutritive meal for the female, even in Mormon crickets. Hemolymph protein was higher in Mormon crickets that mated and consumed the spermatophylax in our study (Table S4, Fig. S3); however, if these females did have higher protein intake, it was not reflected in their microbiome. Although their spermatophylax is relatively large and females compete for spermatophylax-producing males under low nutrient conditions (Gwynne, 1984b, 1993), the nutritional consequences of spermatophylax consumption has not been explicitly measured in Mormon crickets.

Second, nuptial gifts might not influence the gut microbiota because of a lack of sensitivity of the microbiome to dietary protein, irrespective of the nutritional properties of the gift itself. Our experiment supports this hypothesis, as increasing dietary protein did not significantly influence the gut microbiome, at least in decorated crickets (Table S2). Cricket gut microbiomes thus might not confer the same degree of plasticity in resource use as has been suggested for humans (David *et al.*, 2014). Experiments measuring metabolic activity under different dietary regimes are required to test this hypothesis.

Finally, it is possible that spermatophylax consumption could affect the microbiome under a different dietary regime not tested in our study. Mormon crickets in particular occur in habitats that vary widely in available protein and other nutrients (Gwynne, 1984b), and under some conditions in nature spermatophylax consumption might have a greater effect than observed in our experiments.

### Conclusion

Social behavior is emerging as an important factor shaping the diversity of the microbiome (Powell *et al.*, 2014; Smith and Mueller, 2015; Tung *et al.*, 2015; Moeller *et al.*, 2016). Progress in this area requires studies that use experimental manipulations of social interactions to complement surveys that correlate microbiome composition and host traits (e.g. group membership, dominance rank, social interaction networks) to infer their relationship (Archie and Tung, 2015). To our knowledge, our study is the first to use such an experimental approach to demonstrate that sexual interactions affect the structure of the gut microbiome. Given the relative simplicity of their gut microbiomes and their long standing as models in the study of sexual behavior, crickets and katydids provide an exciting opportunity to expand our knowledge of host-microbe symbioses.

## DATA ACCESSIBILITY

Sequences are deposited in Genbank SRA accessions SRP073329 and SRP073374.

## ACKNOWLEDGEMENTS

We thank Laura Senior, Alexis Carlson, Aaron McAughan, Danny Nyugen, Lisa Zezas and Karth Swaminath for help with field collections and performing experiments, Spencer Behmer for the artificial diets and Scott Sakaluk for advice on the decorated cricket methods. This work was funded by US National Science Foundation award DEB-1354666 and the W.M. Wheeler Lost Pines Endowment from the University of Texas at Austin.

